# Structural connectome fingerprinting and age prediction in pediatric development: assessing voxel- and surface-based white matter connectivity

**DOI:** 10.1101/2021.02.19.432024

**Authors:** Noor B Al-Sharif, Etienne St-Onge, Jacob W Vogel, Maxime Descoteaux, Alan C Evans

## Abstract

Mapping structural white matter connectivity is a challenge, with many barriers to accurate representation. Here, we assessed the replicability and reliability of two connectome-generating methods, voxel- or surface-based, using test-retest analyses, fingerprinting and age prediction. The two connectomic methods are initiated by the same state-of-the-art dMRI processing pipeline before diverging at the tractography and connectome-generating steps using either voxels or surfaces. While both methods performed very well across all analyses, voxel-based connectomes performed marginally better than surface-based connectomes. Notably, structural connectomes derived from either method demonstrate reliably accurate representations of both individuals and their chronological age, comparable to similar analyses employing multi-modal features. The difference in methodological performance could be attributed to a number of method-specific features but ultimately show that cutting-edge tractography with robust dMRI processing produces reliable white matter connectivity measures.

## 1. Introduction

The majority of post-natal brain development occurs in white matter (WM), namely through the pruning, refinement or development of axonal connections (Sowell et al., 2004; Lenroot & Giedd, 2006; Giedd & Rapoport, 2010). These microstructural changes underlie larger transformations in both structural and functional whole-brain connectivity (Lebel et al., 2008). Despite advancements in research methods at multiple levels of inquiry, we are still establishing a ‘map’ of how developmental changes manifest anatomically, in complement to well-described cognitive development (Goldman-Rakic, 1987; Nagy et al., 2004; Barrouillet, 2015). Abnormalities in both structural and functional connectivity have been suggested to underlie myriad neurological conditions, disorders and diseases (O’Sullivan et al., 2001; Pievani et al., 2014; Kana et al., 2014). Having a clear, comprehensive reference of how the brain typically looks during development will help to better understand deviations from the norm.

Structural connectivity, a representation of anatomical WM connections, is a measure designed to assess how the brain is internally connected (Jones, 2008; Jbabdi et al., 2015; Sotiropoulos & Zalesky, 2019). Due to a number of methodological limitations, however, characterizing brain connectivity *in vivo* is notoriously difficult (Jones et al., 2013; Jeurissen et al., 2019). Imaging methods typically employed to determine in vivo connectivity, such as T1- or diffusion-weighted magnetic resonance imaging (dMRI), lack the resolution acuity to depict axon-specific connections. Therefore, estimations of structural connectivity are made using diffusion tensor imaging (DTI) such as fractional anisotropy (FA), radial, mean and axial diffusivity (RD, MD, AD), as well as tractography (Basser & Jones, 2002). The latter contributes to one of the most common measures of WM connectivity: streamline count, the number of tractogram streamlines which connect two brain regions. In addition to scanning resolution, the accuracy of these measures is also impeded by a number of issues in tractogram representation such as crossing fibers and the gyral crown bias (Schilling et al., 2018; Girard et al., 2014).

One of the most common methods to measure structural connectivity is the voxel-based approach, which uses a predefined atlas in 3D voxel space to delineate anatomical regions. Tractograms aligned within the same space represent connections between regions of interest (ROIs) when a streamline crosses a labelled voxel region. Though voxel-wise methods are widespread and fairly simple to execute and interpret, using voxels to represent connectivity does present some anatomical limitations. These include structural distortions introduced by nonlinear registration, poor cortical coverage and bias against complex or short-range streamlines (Jbabdi & Johansen-Berg, 2011; Van Essen et al., 2014). Recent advances in cortical surface modeling have improved our ability to represent anatomical connectivity in ways that cannot be captured as efficiently or accurately using voxel-wise approaches (Smith et al., 2012; St-Onge et al., 2018).

Surface-based connectivity uses cortical mesh surfaces, instead of voxels, as the seeding points for initializing and terminating streamlines (Smith et al., 2012; St-Onge et al., 2018). Enhancing tractography with surfaces attempts to bypass certain disadvantages of traditional tractography, generating more precise and uniform endpoint placement across a surface. By using information from the cortical surface directly in creating the connectome, surface-based connectivity also offers the potential to integrate other grey matter (GM) features such as cortical thickness, GM volume and surface area in connectomic analyses.

A key component of a method’s utility lies in its stability, which is often described in the contexts of reliability and reproducibility. These are quantifiable properties which determine how well a method’s results can be replicated and extrapolated to other datasets. The former illustrates a method’s inherent accuracy while the latter depicts its broader real-world relevance. While methods can be validated through replication in another dataset, we can assess our methods within the same dataset by relying on the use of same-session repeated acquisitions.

Here, to evaluate and compare the reliability and reproducibility of both voxel- and surface-based connectivity methods, we employed three analyses: test-retest similarity/distance measures, fingerprinting and age prediction. Using corresponding pairs of imaging and connectivity data, a similarity/distance metric can be measured to evaluate the similarity between two data files from the same subject. The same metric can be calculated between subjects, allowing the computation of inter- and intra-subject similarity/distance averages and variance for each method. Another practical and popular application of test-retest analysis is fingerprinting (Finn et al., 2015), a method which assesses how well subjects can be identified using only their imaging data. This is determined by how accurately a subject’s scan correlates to itself or to another scan from the same subject, as compared to the rest of the dataset. Fingerprinting allows us to understand both how reliable the results are and also how well measures of structural connectivity represent unique individuals. Thirdly, we use age prediction as a tool to validate the biological relevance and reliability of structural connectivity methods. Determining how well a subject’s age can be predicted using structural connectivity illustrates whether such measures accurately and reliably represent normative age across development (Cole et al., 2017; Dlaz-Arteche & Rakesh, 2020). While using white matter-derived features in such data-driven analyses is a budding field, few studies have focused specifically on the use of white matter in age prediction during pediatric development (Brown et al., 2012; Han et al., 2014; Erus et al., 2015; Niu et al., 2020). We execute the test-retest and fingerprinting analyses using same-session test-retest data for 560 subjects from a pediatric imaging dataset and the age prediction in 813 subjects from the same study. These data are preprocessed with *Tractoflow-pve* (Al-Sharif et al., 2020; Theaud et al., 2020), a state-of-the-art diffusion processing pipeline from the Sherbrooke Connectivity Imaging Lab (SCIL), after which the connectomic mapping is completed using two different strategies: voxel- or surface-based. Due to the surface-based method’s more nuanced tractogram construction, we hypothesized that its connectivity representations should provide improved results over voxel-based measures in accurately depicting structural connectivity.

## 2. Methods

Anatomical MRI data were processed initially through one common dMRI processing pipeline, *Tractoflow-pve*, before subsequent processing to generate surface- and voxel-based connectomes. Processing steps are briefly described below; full details are described in (Al-Sharif et al., 2020), which also provides links to the data and processed derivatives.

### 2.1. Participants

Participants, aged 3-22 years, were recruited as part of the Pediatric Imaging Neurocognition and Genetics (PING) study in nine major metropolises across the United States (Jernigan et al., 2016). Here, 821 subjects’ data were used in total (426 male), 560 of which had at least two same session acquisitions for use in the test-retest analyses. In the test-retest group, 8 subjects were missing complete demographic information including age but were retained as these features were irrelevant to the analyses. These subjects were excluded from the age prediction analyses, bringing the total sample for that analysis to 813 subjects. Of these 813 subjects, the mean age was 12.75 years (SD = 4.85 years) and in the test-retest group, the mean age for the 552 subjects with complete demographic data was 13.35 years (SD = 4.61 years). Further details are provided in Jernigan et al. (2016).

### 2.2. MR acquisition

As part of the PING study data collection, multi-modal images were acquired using harmonized protocols on four 3T MR scanners (Siemens Trio-Tim, GE Discovery, GE Signa, Philips Achieva) at nine sites. T1w scans included a motion-reduction protocol to minimize motion artefacts commonly found in pediatric datasets. DMRI acquisitions consisted of 32-direction scans, including 2 baseline b0 volumes, with a single shell b-value of 1000. From PING, we initially gained access to a total of 954 subjects with both T1w and dMRI volumes, of which 634 had at least two or more repeated same-session scans. Subjects were quality assessed at multiple steps of processing and some were excluded for major scanning artefacts, incomplete demographic information or failed preprocessing. In total, 1543 unique acquisitions were used in this study, from 821 subjects, 560 of which had multiple same-session scans.

### 2.3. T1w and dMRI processing

*Tractoflow* (Theaud et al., 2020), developed by SCIL, is a state-of-the-art dMRI processing pipeline which takes raw dMRI, complementary T1w MRI and bvec/bval files as input, and generates processed diffusion metric derivatives including tractography. For our purposes, we modified the original *Tractoflow* pipeline to include the whole-brain mask and partial volume estimation (PVE) maps for GM, WM, CSF and subcortical GM from CIVET 2.1 (Ad-Dab’bagh et al., 2006) along with the initial dMRI input. The pipeline is openly available through the NIMH Data Archive and GitHub as Tractoflow-pve (Al-Sharif et al., 2020).

Through *Tractoflow-pve*, T1w images are processed using their given corresponding CIVET 2.1-produced brain masks and PVE segmentation files. T1w images are masked, de-noised and N4 bias-field corrected before undergoing nonlinear registration to the diffusion image. Their complementary PVE and binary mask files also undergo the same image registration. Diffusion-weighted volumes and corresponding b0 images files are masked, denoised, bias-field and topup-eddy corrected along with their bvec and bval files. Diffusion tensor imaging (DTI) metrics are extracted and computed from the dMRI volume to generate FA, RD, AD, and MD image volumes, as well as fibre orientation distribution function (fODF) volumes.

### 2.4. Tractography

Within *Tractoflow-pve*, structural tractograms were generated using Particle Filtering Tractography (PFT) (Girard et al., 2014), which employs WM-GM-CSF tissue maps to demarcate “include” and “exclude” regions of the brain. “Include” regions define areas of WM where valid, anatomical stream-lines are likely to reach. The resulting tractograms are an approximate representation of WM macrostructure and connections within the brain. A subsequent processing pipeline called *Surface-Enhanced Tractography (SET)* (St-Onge et al., 2018), was employed to generate surface-based tractograms and structural connectomes. An adaptation of the PFT algorithm, SET uses surfaces instead of tissue maps to create streamlines initalized at the GM-WM boundary of cortical and subcortical surfaces. Cortical surfaces are output by CIVET 2.1 directly whereas subcortical surfaces, including the brainstem, were converted from CIVET 2.1 ANIMAL segmentation labels (Collins et al., 1999) using a marching cube algorithm. SET combines these surfaces with dMRI output from *Tractoflow-pve* to reconstruct WM connectivity.

### 2.5. Template construction

To avoid using an adult brain template for this pediatric dataset, a study-specific common space T1w template was created using Advanced Normalization Tools’ (ANTs) iterative multi-variate template construction script (Avants et al., 2011). The resulting transformation files were used to register the voxel-wise DKT atlas into native subject T1w and dMRI space to generate the voxel-based connectivity matrices, as described below.

### 2.6. Connectomes

Voxel-based connectomic matrices were generated using a Desikan-Killiany-Tourville (DKT) (Desikan et al., 2006) segmentation label aligned to each subject’s native diffusion space. To ensure the same subcortical regions were used in both surface- and voxel-based methods, the DKT cortical atlas was combined with subcortical labels from CIVET 2.1 ANIMAL segmentation in each subject’s T1w space before registration to diffusion space. The dMRI-registered segmentation label and the subject’s tractogram were used to create acquisition-specific connectivity matrices. Connectivity is represented as the number of tractography streamlines with endpoints within a ROI as delineated by the DKT label. Surface-based connectomic matrices were generated using SET by counting the number of streamlines which intersect two cortical or subcortical regions of a surface. Surfaces are delineated into ROIs using cortical labels in the CIVET DKT surface atlas. Connectomes were subsequently normalized by the sum of their total streamline count to facilitate comparison between methods.

### 2.7. Analysis

#### 2.7.1. Test-retest

To assess the variability of our reconstructed connectivity matrices, we computed the distance between two acquisitions from the same subject (inter-subject) and other subjects (intra-subject). This metric informs how similar two matrices are to each other and also how distinguishable they are from matrices of a different subject. For both approaches, we calculated the mean L1 distance for both inter- and intra-subject comparisons and determined the ratio of inter/intra distance (Aggarwal et al., 2001; Allen et al., 2014; St-Onge et al., 2018). A lower L1 distance indicates more similarity between two connectomes, where a distance of 0 would represent two identical matrices. Thus, a higher ratio indicates a better ability both to match subjects to themselves (intra L1) and also distinguish them from others (inter L1). The L1 distance between two connectivity matrices *(A, B)* is defined by the sum of the absolute difference between each connection weight (c), ∑∀_c_ |A_c_ - *B_c_*|. Results from each method were compared using a t-test.

#### 2.7.2. Fingerprinting

In addition to the above assessments, we used data from the 560 subjects with multiple same-session scans to implement a fingerprinting analysis. The first and second scans of all subjects with at least two acquisitions were extracted as follows into two separate sets of data. First, each connectome’s diagonal was set to zeros before symmetrizing and normalizing the matrices. Symmetrized matrices are created by averaging the connectivity between corresponding pairwise connections, and matrices were normalized by dividing each connection by the sum of all pairwise connections within the matrix. Data from each connectome were then vectorized, or flattened, and appended to a 2D matrix for either “first” or “second” scans. To determine the similarity between a “first” scan and all other “second”, or repeated, scans, we ran a Pearson’s correlation between each acquisition’s connectome and the connectomes of every other repeated acquisition in the dataset. Fingerprinting is considered successful if the maximum correlation value results from the comparison of two scans from the same subject. The fingerprinting accuracy between the voxel- and surface-based methods was then compared with a χ^2^ test.

#### 2.7.3. Age prediction

While the previous approaches assess and compare the reproducibility of the two diffusion pipelines, we also wished to assess and compare how well data prepared using these methods performed in real-world applications. We chose to submit these data to a machine learning pipeline designed to predict participant age using purely neuroimaging information. Age prediction was chosen as it is a fairly easy problem with wide application across the neuroimaging field (Cole et al., 2017; F da Costa et al., 2020; Niu et al., 2020; Han et al., 2014; Tønnesen et al., 2020; Liem et al., 2017; Erus et al., 2015; Díaz-Arteche & Rakesh, 2020), and because it is relevant to the biology of neurodevelopment. Our goal was to establish the maximum reliable age prediction accuracy (and minimum age prediction error) using basic machine learning approaches and the preprocessed imaging data as input. The machine learning pipeline was applied separately for datasets preprocessed using voxel-wise and surface-based approaches, and output from the best model from each dataset were compared. We were also interested in whether additionally adding cortical thickness data could improve predictions.

As above, the diagonal of each subject’s tractographic connectivity matrix was set to 0s, the matrices were symmetrized and normalized. For subjects with more than one same-session scan, matrices derived from all acquisitions were averaged such that each subject had only one connectivity matrix. The lower triangle of each matrix was then extracted, flattened, and appended to one another, creating a 2D subject x (connectivity) feature matrix. A similar subject x feature matrix was created for cortical thickness (CTx) by appending vectors for each subject containing the cortical thickness of each brain region. Both types of feature matrices were normalized using a standard scaler. To partially account for differences in scanner models, we included dummy coded variables for each of the scanner models into the feature matrices. Finally, we set aside data from 20% of the participants as a final “lock-box set”, and ensured that the proportion of subjects younger than 8, 8-12, and 13+ was equal between the lock-box set and the rest of the data.

The following analysis was conducted using the python package scikitlearn (Pedregosa et al., 2011). We next built a grid search that cycled through different combinations of pipeline steps, machine learning estimators, imaging features and hyperparameters. Specifically, the following pipeline aspects were varied:

- Input features: WM connectivity only, CTx only, WM and CTx features
- Feature dimension reduction: None, PCA (100 components), only features with a significant (FDR p<0.05) correlation with age (f_regression)
- Estimators and hyperparameters:

– LassoCV
– Support vector regressor (SVR) w/ Linear kernel w/ C chosen from the array[0.0001,0.001,0.1,1,10]
– Random forest regressor (RFR) w/ max depth selected from the array [3, 5, 7, 10, 15, 20, 40, 50, None]

Hyperparameters for all models were chosen using a nested cross-validated grid search (GridSearchCV), except for Lasso, which has a built-in cross-validation framework for determining the value for the alpha parameter. All possible combinations of these pipeline inputs were modeled with the exception that no PCA was applied for models using CTx only features as the number of components exceeded the number of input features.

For each pipeline, thirty 80/20 training/validation splits were created from the data. For each split, the Pipeline was fit to the training data and used to predict the age of unseen individuals in the testing data. The mean squared error (MSE) and R^2^ score were recorded for each split in order to establish a range of uncertainty within the predictions of each pipeline. For both the voxel-wise and SET processed data separately, the best performing model was selected based on having the lowest MSE and/or highest R^2^ score. In the event of more than one model with equivalent scores, the model with the lowest number of features was selected. Finally, the best performing model was evaluated on the left out 20% lock-box set, and this performance was compared across the two WM preprocessing pipelines. To validate that the selected model’s performance was not influenced by the specific lock-box sample, 100 new train-test splits were generated from the total dataset and trained using the only selected model. The distribution of R^2^ and MSE results on the new 100 test sets were then compared between both voxel and surface methods. To assess whether subcortical regions were driving prediction results, i.e. regions in which SET might be less effective, the analyses were repeated using only cortical ROIs, again comparing R^2^ and MSE results.

#### 2.7.4. Resampling

To test whether our results were robust to arbitrary decisions in stream-line seeding count, we resampled all connectomes for each method at various counts of resulting streamlines. This allowed us to evaluate whether the test-retest variability of a method would change over a range of total streamline counts. The original voxel-based connectomes were initialized with a total of 50 million (M) seeds while the surface-based connectomes totalled 25M seeds. However, the resulting amount of reconstructed streamlines varies per subject and averaged to *5M for PFT and *10M for SET. For this reason, resulting connectomes were resampled at specific total streamline counts: 100k, 250k, 500k, 1M, 1.5M, and 2M. In addition, to resample the connectomes by seed count, the number of streamline count was weighted by the percentage of seeds resulting in a streamline. This resampling was done at 100k, 250k, 500k, 1M, 1.5M, 2M, 3M, 5M, 7.5M, 10M, and 15M total seeds. We then re-ran the fingerprinting and age prediction analyses with the resampled connectomes, grouped by preprocessing method (voxel or surface), resampling reference (seeds or streamlines), and matrix resolution (seed or streamline count), and compared the results between corresponding groups of voxel and surface connectomes.

## 3. Results

### 3.1. Test-retest

Test-retest L1 distance (Table 1) for voxel-derived connectomes showed an intra-subject distance average = 0.144 (0.061 SD) that was significantly lower (T = −73.4, p < 0.0001) than that of surface-based connectomes at 0.231 (0.061 SD). Alternatively, voxel connectomes displayed an inter-subject distance of 0.418 (0.021 SD) which was also lower than that of the surface connectomes at 0.554 (0.032 SD). The subsequent intra/inter distance ratio for voxel connectomes was 2.900 and that of the surface connectomes was 2.394. Similar results were obtained using L2 and χ^2^ distance.

**Table 1:**
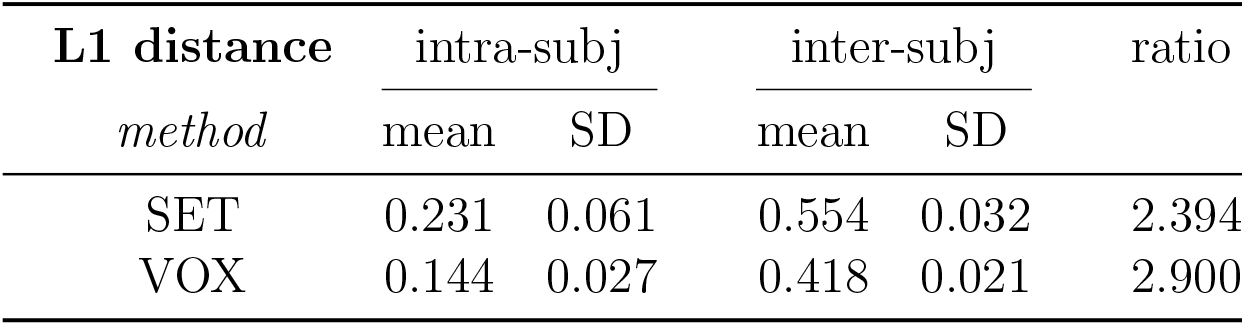
L1 distance metric results where intra-subject distance measures similarity of data from same subject and inter-subject distance measures similarity between data from different subjects. Intra/inter ratio indicates better ability to self-identify and distinguish from others.

| L1 distance<br>*method* | intra-subj |  | inter-subj |  | ratio |
| --- | --- | --- | --- | --- | --- |
|  | mean | SD | mean | SD |  |
| SET | 0.231 | 0.061 | 0.554 | 0.032 | 2.394 |
| VOX | 0.144 | 0.027 | 0.418 | 0.021 | 2.900 |

### 3.2. Fingerprinting

Fingerprinting analyses using voxel-based connectomes were able to match subjects to their own scans with 99.8% accuracy, correctly identifying 559 subjects out of 560 (Figure 1). This performance was significantly better (χ = 29.6, p < 0.0001) than that of surface-based connectome fingerprinting showed 94.6% accuracy, correctly identifying 530 subjects.

**Figure 1:**
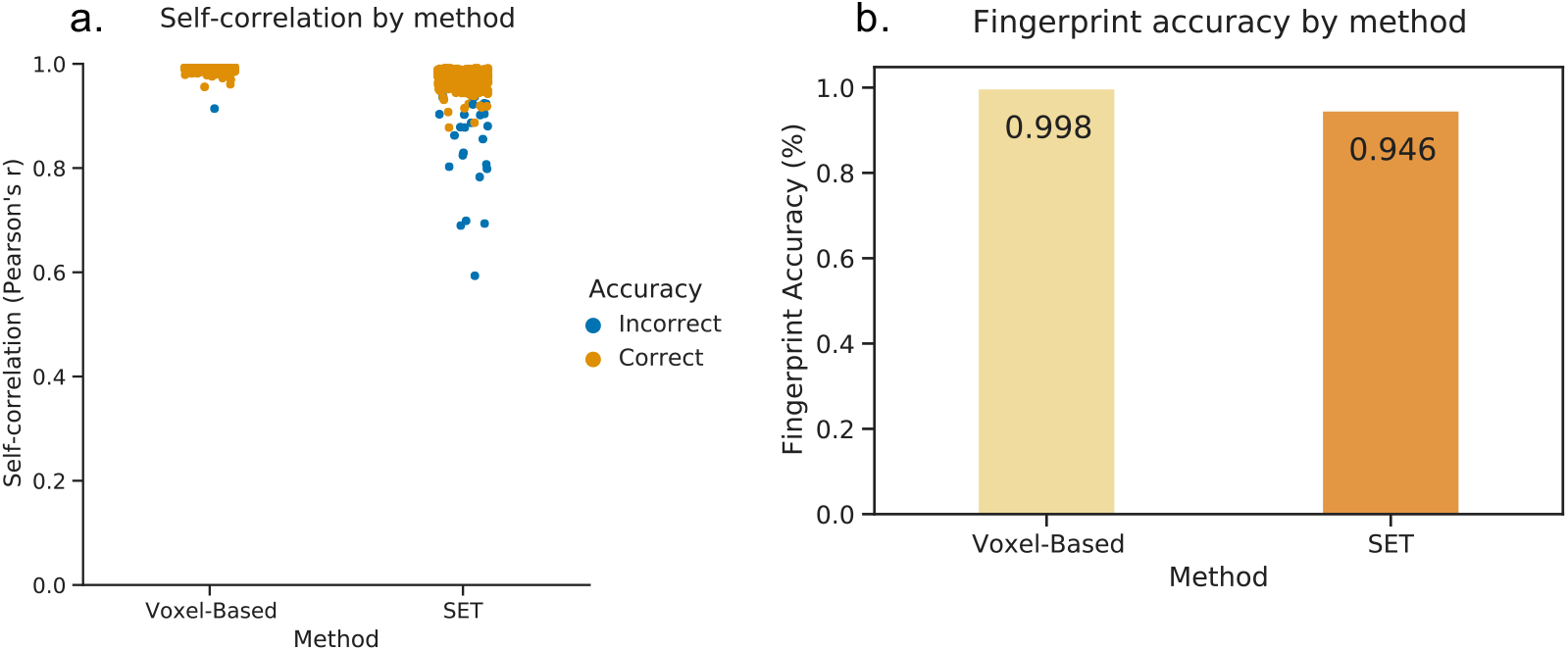
**a.** Correlation of each subject to itself for both connectomic methods, determined by a Pearson’s r. Subjects whose highest correlation resulted from a correct self-identification are marked in orange, incorrect instances are in blue. **b.** Fingerprint accuracy per method, as a percent of correctly identified matches.

### 3.3. Age prediction

After iteratively training and testing multiple models with 80% of the total dataset, we evaluated the average test-set R^2^ and MSE of each model to determine the best fit for both connectomic methods (Figure 2). We also prioritised using models with fewer features. The best fitting model using voxel-based connectomes was Lasso Select using both GM (CTx) and WM (connectivity) features, with an average R^2^ of 0.804 (range = 0.728-0.833, mean MSE = 4.48, Figure 2a and c). Using CTx features alone resulted in an average R^2^ of 0.711 (0.635-0.765) and using WM connectivity alone resulted in an average R^2^ of 0.733 (0.637-0.797). We then applied this model to the final left out 20% lock-box set for voxel-based connectomes, resulting in an R^2^ of 0.824 (MSE = 4.46, MAE = 2.11, Figure 2e). Feature weights for which voxel-wise connections contributed most to the age prediction model are depicted in Figure 3a.

**Figure 2:**
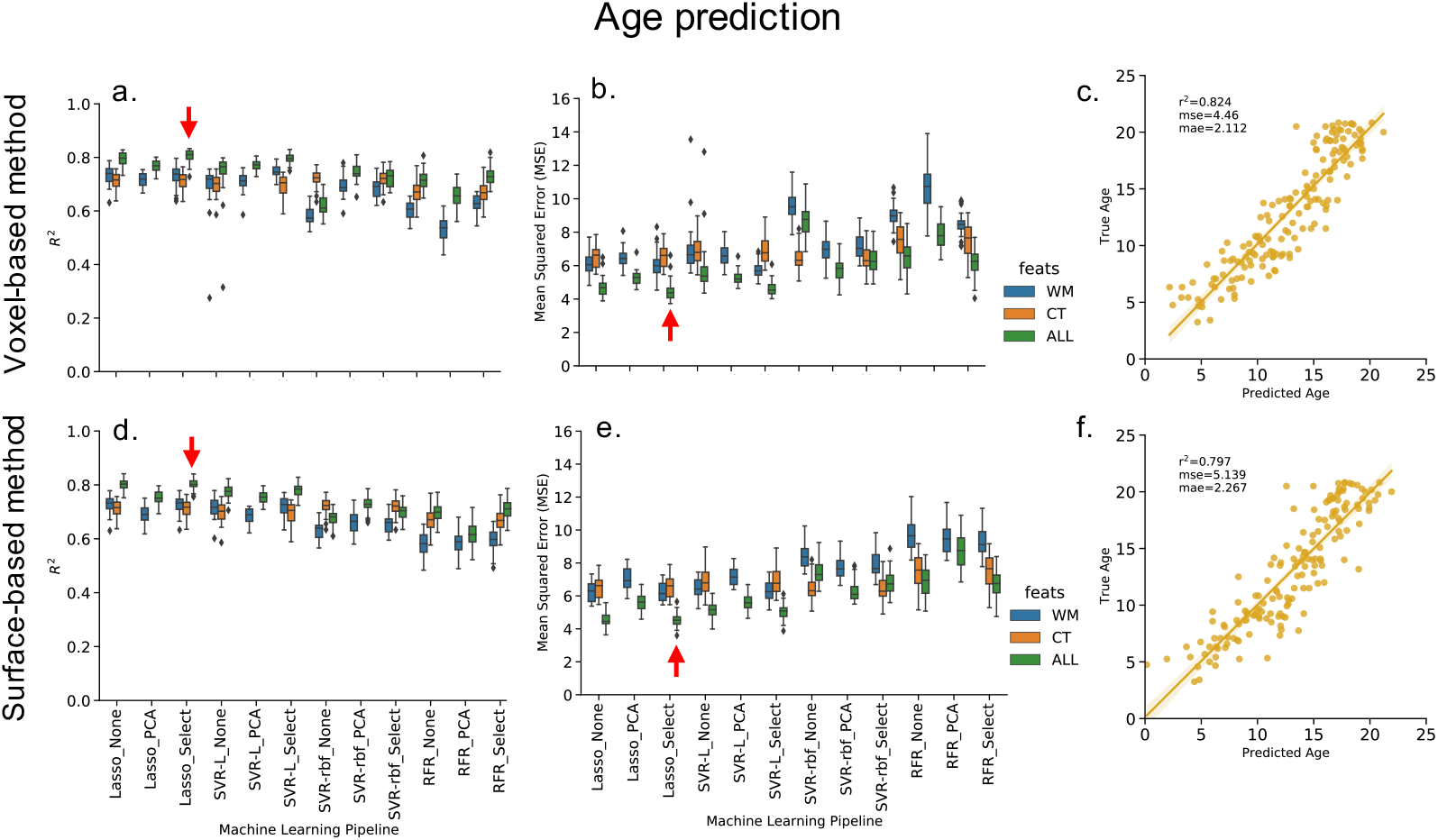
Average results for all models trained and tested using connectomes from both methods: a and b show results using voxel-based connectomes; d and e show results using surface-based connectomes. Each machine learning pipeline was tested using WM connectivity only (blue), CT only (orange) and ALL features (green). Pipeline with the highest average R^2^ and lowest average MSE when applied to test set (indicated by red arrows) was selected as model to evaluate on lock-box data (c and f).

**Figure 3:**
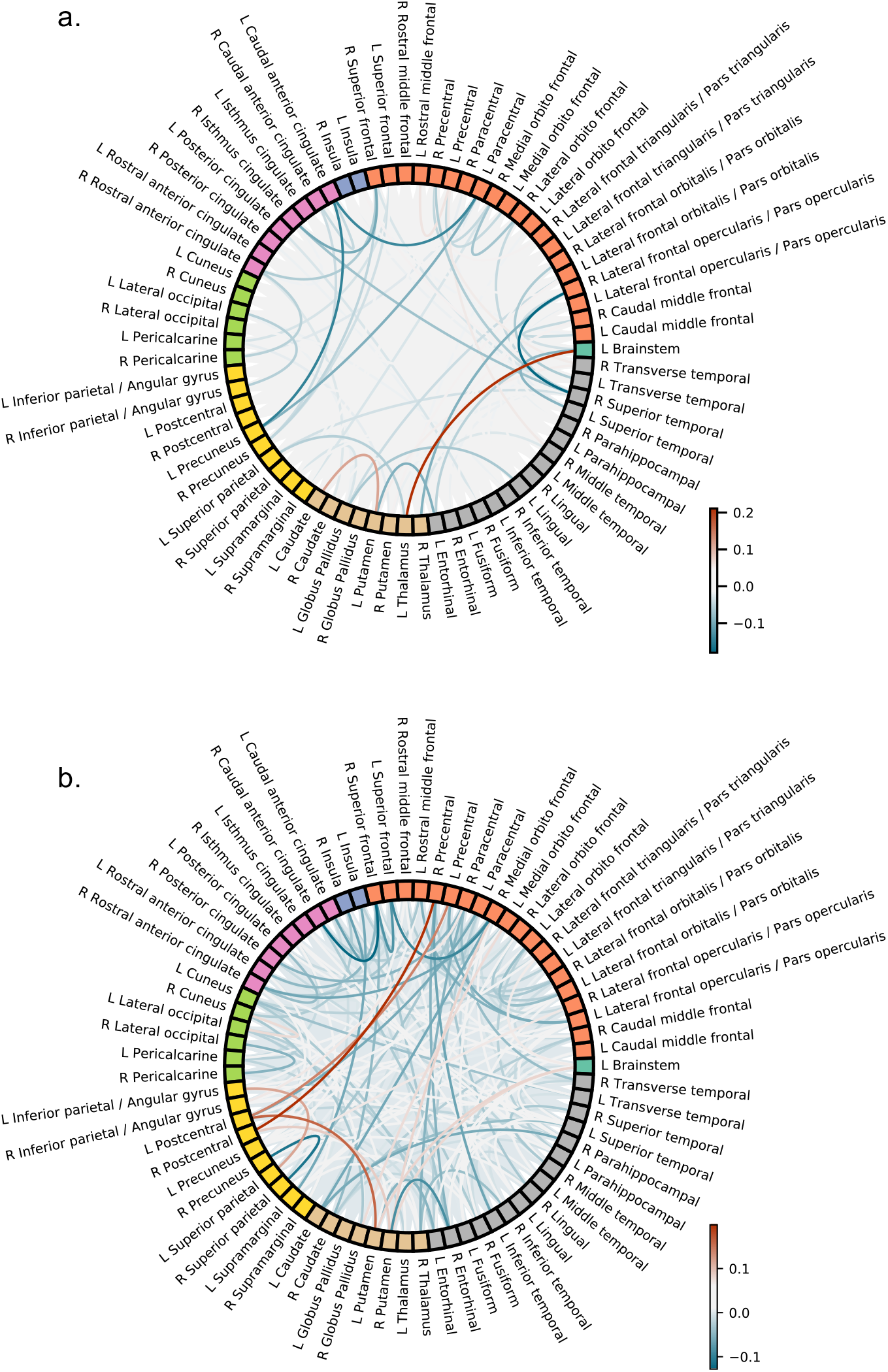
Feature weights: connectivity features which contributed most strongly to the age prediction model for a. voxel-based connectomes and b. surface-based connectomes, depicted as an increase with age within that connection in red and a decrease in blue.

The best fitting model for surface-based connectomes was also Lasso Select using both GM and WM features with a mean R^2^ of 0.802 (range = 0.756-0.840, mean MSE = 4.52) (Figure 2b and d). Using GM alone resulted in an average R^2^ of 0.711 (range = 0.635-0.765), while using WM connectivity alone resulted in an average R^2^ of 0.726 (range = 0.633-0.779). In the final lock-box set, this model resulted in an R^2^ of 0.797 (MSE = 5.14, MAE = 2.27) (Figure 2f). Feature weights for which surface-wise connections contributed most to the age prediction model are depicted in Figure 3b.

R^2^ and MSE distributions from the 100 train-test samples using only the best performing model showed that age prediction performance was not significantly different between voxel and surface methods (R^2^: p = 0.42, MSE: p = 0.41). These results were unchanged when using cortical ROIs only.

### 3.4. Resampling

While the accuracy of the fingerprinting analyses varied slightly with resampling, the results from voxel-based connectomes (seeds mean = 0.997, streamlines mean = 0.998) were still consistently higher than those of the surface-based connectomes (seeds mean = 0.943, streamlines mean = 0.945), at all resolutions (Figure 4, Table 2). Age prediction results also varied little from the full resolution connectomes, with voxel-based connectome predictions (seeds mean R^2^ = 0.814, streamlines mean R^2^ = 0.811) almost exactly equal to those of the surface-based connectomes (seeds mean R^2^ = 0.810, streamlines mean R^2^ = 0.810), regardless of the seed or streamline resolution.

**Figure 4:**
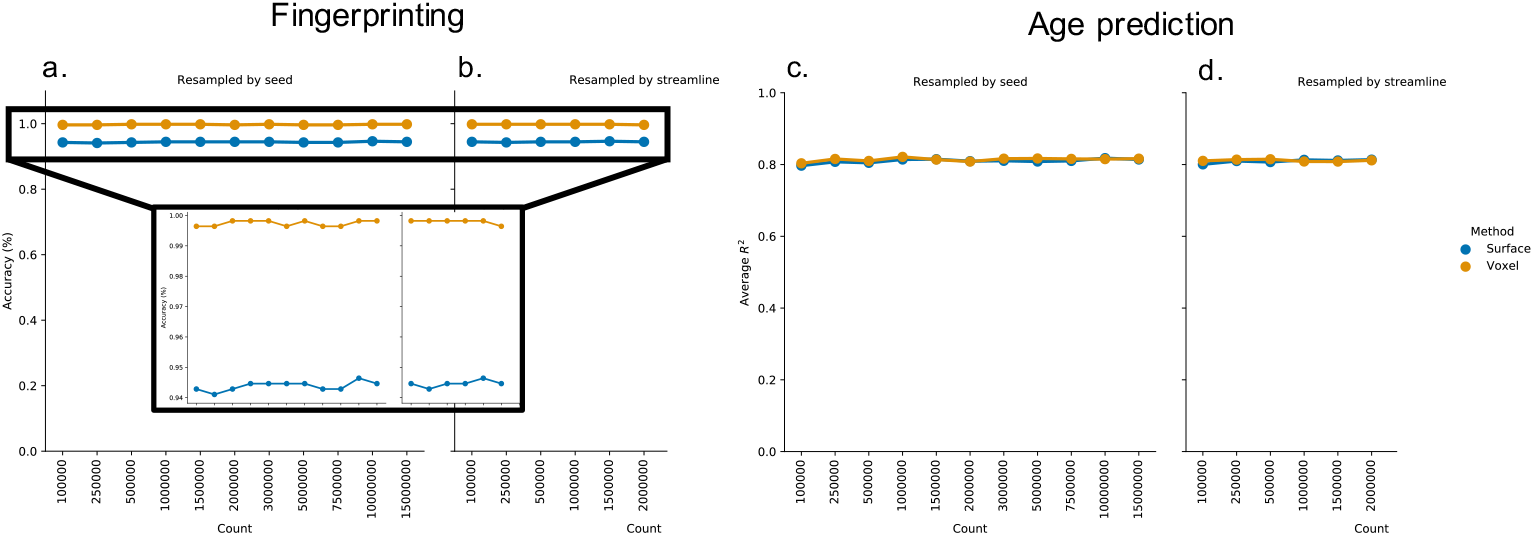
Mean fingerprinting accuracy and age prediction R^2^ results across all resampled connectomes, using either seeds (a,c) or streamlines (b,d) as the resampling reference. Inset shows close-up of differences in fingerprinting accuracy between methods.

**Table 2:** Average age prediction (R^2^) and fingerprinting (accuracy) results for resampled connectomes, grouped by resampling reference (seeds or streamlines) per connectomic method.

| Seeds<br>*method* | $R^2$ | | Accuracy (%) | |
| --- | --- | --- | --- | --- |
|  | mean | range | mean | range |
| VOX | 0.814 | (0.803, 0.821) | 0.997 | (0.996, 0.998) |
| SET | 0.810 | (0.797, 0.818) | 0.944 | (0.941, 0.946) |
| Streamlines<br>*method* | $R^2$ | | Accuracy (%) | |
|  | mean | range | mean | range |
| VOX | 0.811 | (0.808, 0.815) | 0.998 | (0.996, 0.998) |
| SET | 0.810 | (0.801, 0.814) | 0.945 | (0.943, 0.946) |

## 4. Discussion

The quantification of whole-brain white matter structure presents a process rife with pitfalls and erroneous estimation. To assess the methods available for overcoming these limitations when measuring structural connectivity, we employed both traditional and more recently developed preprocessing techniques. Despite the potential advancements offered by using Surface-Enhanced Tractography, connectomes from voxel-based pipelines outperformed surface-based connectomes in reproducibility metrics and produced equivalent results in age prediction. While results varied slightly between the two, both pipelines generated connectivity derivatives which were highly reliable and reproducible.

Both pipelines were able to match subjects to their own data with very high accuracy, though using voxel-based connectomes achieved slightly better results than those of surface-based connectomes. The higher L1 inter/intra ratio for voxel-based connectomes indicated that voxel-based connectomes were more easily distinguished solely based on the connectivity matrix similarity. This result was unexpected as we predicted that with more enhanced, biologically accurate streamline coverage provided by SET, subjects would have more nuanced and individualised tractograms and subsequent connectomes. However, SET’s higher frequency streamline improvements may have introduced more noise into the connectomes, increasing the L1 distance, making them harder to distinguish from each other. The enhancements may also have been negated by the use of a broad brain parcellation such as DKT,rendering these improvements inconsequential for this particular analysis. Alternatively, the voxel-based connectomes might be subject to subject-specific deformities introduced by nonlinear registration, which would make an individual more easily matched to their own data. These results outperform the first instance of connectomic fingerprinting using rsfMRI which presented 89% accuracy using whole-brain connectomes (Finn et al., 2015). Parker et al. (2016) first showed that anatomical white matter data can be used as fingerprints based on tract shape, with 93% accuracy, which both our connectome groups surpassed. Another study of the Human Connectome Project (Kumar et al., 2018) compared imaging data from multiple modalities and found that, while using FA from dMRI showed better fingerprinting capabilities than T1w features alone, a combination multi-modal features offered the best “separability”. Here, the voxel-based connectomes outperformed any of the dMRI metrics used on their own in Kumar et al. (2018) and matched the results using multi-modal combinations. White matter features used in fingerprinting may outperform resting-state connectivity due to their reliance on defined anatomical structure rendering better reliability between scans, as opposed to state-dependent connectomes. Notably, regardless of preprocessing pipeline, differences in individual structural connections as represented by tractography are measurable to the extent that they can be used reliably for unique individual identification.

While the age prediction results on the lock-box set was qualitatively higher using voxel-based connectomes, the range of results from an iterative sample show that this difference is not significant. Given the wide age range of the PING dataset, the defining drivers of age prediction may lie in changes within more principal, long-range connectivity patterns, such as those that transverse the subcortical structures, and not in areas of nuanced local connectivity, which are more likely to have been enhanced by SET. Subject specificity to such an individualized degree at the local level would be irrelevant if the larger effects of age were determined by other regions or measures. When subcortical ROIs were removed from the analysis, however, the results remained as is. This might suggest these results are similarly impacted by SET-induced high frequency noise or DKT-induced flattening of high frequency data, as in the fingerprinting. Interestingly, the plotted weights of connectivity features which contributed to the prediction model differ widely between voxel- and surface-based methods (Fig. 3). Voxel-based prediction models show sparse feature plots, with few regions contributing to the model, whereas the surface-enhanced connectivity plots are far denser, likely due to SET’s on-average increased streamline count and cortical coverage across the whole brain. However, this may also indicate SET’s ability to incorporate more whole-brain connectivity features into its connectomic representation as more connections are generated in SET’s tractograms. SET might improve modeling and representation of other connectivity measures in other contexts, especially when coupled with grey matter features, but these potential improvements could be far too small to have an impact in our current analyses. SET may also introduce other sources of error that prevented it from outperforming voxel-based methods. Other studies in pediatric or lifespan age prediction have shown comparable results, the best of which consistently use multi-modal models including more features than we have employed here (Brown et al., 2012; Erus et al., 2015; Liem et al., 2017; Han et al., 2014; Niu et al., 2020; Cole et al., 2017). However, we do show that even with fairly basic preprocessing tools and limited measures, high age prediction accuracy can still be achieved.

Hypothesizing that SET’s advanced tractogram modeling might outperform voxel-based methods with fewer streamlines, i.e. at lower sampled connectomes, we resampled all connectomes at varying levels of seed and streamline count. Resampling had almost no effect on either the fingerprinting or age prediction results. Fingerprinting accuracy using voxel-based connectomes was nearly identical at every instance of resampling by streamlines. Marginally less consistency is seen when resampling by seeding point, though the range in accuracy is the same as in streamline resampling; streamline resampling appeared to be most consistent using between 500k and 1.5m streamlines. Surface-based connectomes also show stable results regardless of resampling, with the most consistent results when resampling by seeds between 1m and 3m seeding points. In either case, these results suggest that highly accurate identification of individual subjects using structural connectomes can be consistently achieved with an average minimum of 1m seeding points, forgoing the need for more computationally demanding processing at higher resolutions. However, the potential increased connectivity and higher frequency impacts of SET may not be captured in our current analysis.

## 5. Conclusion

In summary, our results indicate that structural connectomes, regardless of methodology, can be used reliably for subject-specific identification and capture biologically relevant characteristics of development. Ultimately, however, the strength of these connectomes may lie in the dMRI processing pipeline that initiates both voxel- and surface-based methods. With a robust, reliable processing pipeline such as *Tractoflow*, resulting tractograms lead to reliable and informative structural connectomes. Using voxel-based atlases to create connectomes is a common and quick technique that may be sufficient for high-level analyses. Surface-based methods will take more time and computational resources, but may be justifiable if subsequent analyses call for integrating grey matter or other surface-based features such as cortical thickness.

## Data Availability Statement

The data that support the findings of this study are available in NIMH Data Archive (NDA) at 10.15154/1519178, study ID #932 ([dataset] Al-Sharif et al., 2020). These data were derived from the following resources also available in NDA as Pediatric Imaging, Neurocognition, and Genetics (PING) study ID #2607.

## Acknowledgements

Supported by grants from Brain Canada (238990, 243030), CFREF/HBHL Innovative Ideas (247613), Coutu Research Fund (241177), CFREF/HBHL Discovery (247712), Universitíe de Sherbrooke Research Chair in NeuroInformatics and NSERC Discovery Grant.

